# How Cognitive and Reactive Fear Circuits Optimize Escape Decisions in Humans

**DOI:** 10.1101/207936

**Authors:** Song. Qi, Demis. Hassabis, Jiayin Sun, Fangjian. Guo, Nathaniel Daw, Dean Mobbs

## Abstract

Flight initiation distance (FID), the distance at which an organism flees from an approaching threat, is an ecological metric of cost-benefit functions of escape decisions. We adapted the FID paradigm to investigate how fast or slow attacking ‘virtual predators’ constrain escape decisions. We show that rapid escape decisions rely on ‘reactive fear’ circuits in the periaqueductal gray and midcingulate cortex (MCC), while protracted escape decisions, defined by larger buffer zones, were associated with ‘cognitive fear’ circuits which include posterior cingulate cortex, hippocampus and the ventromedial prefrontal cortex, circuits implicated in strategic avoidance and behavioral flexibility. Using a Bayesian Decision Model, we further show that optimization of escape decisions under rapid flight were localized to the MCC, a region involved in adaptive motor control, while the hippocampus is implicated in optimizing decisions that update and control slower escape initiation. These results demonstrate an unexplored link between defensive survival circuits and their role in adaptive escape decisions.

**Significance:** Humans, like other animals, have evolved a set of circuits whose primary function is survival. In the case of predation, these circuits include ‘reactive fear’ circuits involved in fast and immediate escape decisions and ‘cognitive fear’ circuits that are involved in the conscious feeling of threat as well as slow strategic escape. Using neuroimaging combined with computational modeling, we support this differentiation of fear circuits by showing that fast escape decisions are elicited by the periaqueductal gray and MCC, regions involved in reactive flight. Conversely, slower escape decisions rely on the hippocampus, posterior cingulate cortex and prefrontal cortex, a circuit implicated in behavioral flexibility. These results support the role of the defensive survival circuitry in escape decisions and a separation of fear into reactive and cognitive circuits.

## Introduction

Survival depends on the adaptive capacity to balance fitness promoting behaviors such as copulation and foraging with the omnipresent risk of lethal predatory attack (Cooper and Blumstein, 2015; Mobbs and Kim, 2015). In the field of behavioral ecology, this balance between survival behaviors is depicted by economic models of flight initiation distance (FID) that capture risk functions by measuring the distance at which an organism flees from an approaching threat, while considering the cost of fleeing (Ydenberg and Dill, 1986; 2015). A wealth of ethological literature demonstrates that prey are remarkably adept at escape and make decisions based on the predator’s directionality, lethality, velocity, and prior encounters with the predator (Stankowich and Blumstein, 2005). In addition to its capacity to measure adaptive behavior, FID is a well-established index of threat sensitivity resulting in large variability within and between species. Despite being applied to a large variety of taxa, this reliable measure has not been used to identify heterogeneity in threat sensitivity or escape decisions in humans, and the neural circuits remain unexplored.

Theoretical and neuroanatomical models support the existence of an interconnected defensive survival circuitry that is remarkably preserved across species (Blanchard and Blanchard, 1990; Panksepp, 1998, 2011; Price, 2005). Under the conditions of immediate danger, the ‘reactive fear’ circuitry is evoked midbrain PAG, central amygdala (CeA), hypothalamus and the midcingulate cortex (MCC) relay, update and initiate essentially innate reactions including motor responses such as flight and freezing (Panksepp, 2011; Shackman et al., 2011; Gross and Canteras, 2012; Tovote et al., 2016). Conversely, the ventromedial prefrontal cortex (vmPFC), posterior cingulate cortex (PCC), hippocampus and basolateral amygdala, a collective set of regions that constitute the ‘cognitive fear” circuitry, promote behavioral flexibility, strategic decision-making and avoidance (McNaughton and Corr, 2004; Mobbs and Kim, 2015; Price, 2005; LeDoux and Pine, 2016). Although few behavioral ecologists have considered the neurophysiology underlying escape decisions, some have proposed similar dichotomies suggesting that fast, but inaccurate decisions are processed by subcortical regions, while slow, but accurate decisions are processed by cortical system (Trimmer et al., 2008). Cognitive and reactive fear circuits work in harmony by adaptively switching between survival circuits to engage the most optimal strategy to maximize survival (LeDoux, 2012; Davis et al., 2010; Mobbs et al., 2015).

Excitation and inhibition between these circuits is determined by the spatiotemporal distance to the threat (Blanchard and Blanchard, 1990; Halladay and Blair, 2015; McNaughton and Corr, 2004; Mobbs et al., 2015). For example, distant threat often results in freezing and threat assessment, yet when the threat is close active flight and fight are observed. Distance to the threat, therefore, is crucial in choosing the best escape strategy. Evidence suggests that this is conserved across species. In humans, active escape tasks have been used, where the goal of the subject is to escape from a virtual looming threat – a red dot with the capacity to chase, capture and shock the subject in a virtual maze. Functional MRI results show that when a threat is distant, there is increased activity in the vmPFC, PCC and BLA. However, as the threat moves closer, there is a switch to increased activity in the central amygdala (CeA), MCC and PAG (Mobbs et al., 2007, 2010). These, and related studies, however, have failed to investigate the neural basis of escape decisions (i.e. flight initiation) or exam the computational mechanisms that underlie escape decisions to changing attack distances.

We developed a paradigm from which to investigate how the defensive survival circuitry facilitates escape decisions when subjects encounter fast or slow attacking threats (Fig 1A). In this task, participants encountered virtual predators of three colors, each representing different attack distances (AD). On each trial, the actual AD was drawn from a Gaussian distribution that was unique to the particular predator type. Fast AD predators (i.e. far or early attacking) were characterized by the virtual predator quickly switching from slow approach to fast attack velocity, therefore requiring the subject to make quick escape decisions. On the other hand, slow AD predators (i.e. close or late attacking) slowly approached for longer time periods resulting in larger buffer zones leading to more time to strategize escape. (Note that “fast” and “slow” here describe the timing of the predator attack, not the speed of the predators) The goal of the task is to try and successfully escape, while at the same time, attempt to acquire as much money as possible by fleeing as late as possible (Fig. 1B). Using this task, we propose several hypotheses: (i) for fast escape decisions, we expect to see activity in the ‘Reactive Fear’ circuitry, while slow escape decisions will reveal more pronounced activity in the ‘Cognitive Fear’ circuitry; (ii) using a Bayesian decision model where subjects’ preference to reward and avoidance to punishment are considered, we predict that the fear and anxiety circuitry will play a role in facilitating fast and slow escape decisions, respectively.

**Fig. 1.**
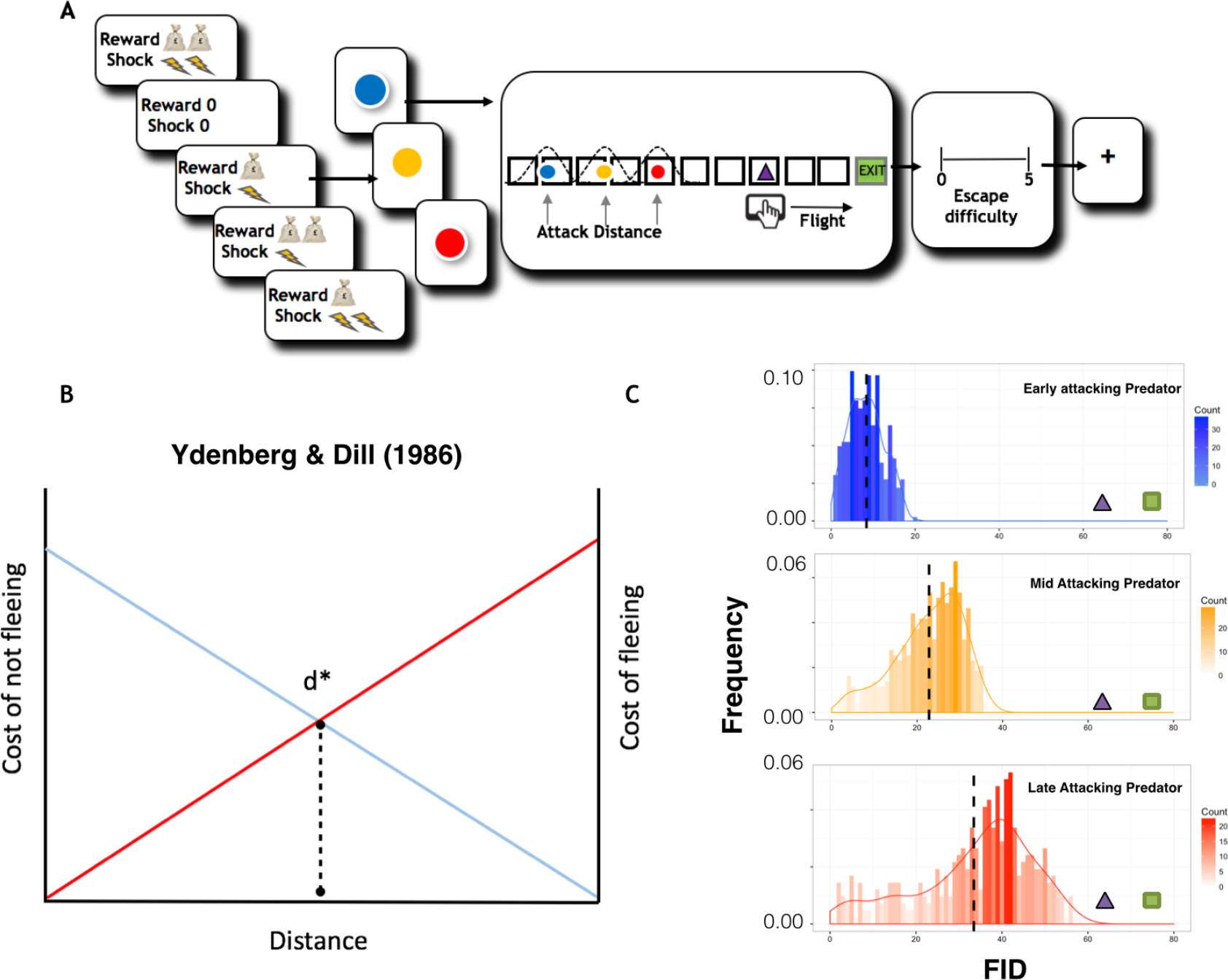
Experimental procedures, Ydenberg and Dill model and distribution of escape decisions. A): Subjects are told whether their decisions will result in high or low reward or shock. They are then, presented with the image of the virtual predator where the color signals the attack distance (2s). (e.g. blue = fast, red = slow). After a short interval, the virtual predator appears at the end of the runway and slowly move towards the subject’s triangle. After an unspecified amount of time (e.g. 4-10s) the artificial predator will attack the subject’s virtual triangle exit (i.e. attack distance). To escape, the subject must flee before the predator attacks. If the subject is caught, they will receive a tolerable, yet aversive, shock to the back of the hand. Trials end when the predator reaches the subject or the exit. To motivate longer fleeing time, the task will include an economic manipulation, where subjects will obtain more money the longer they stay in the starting position and lose money the earlier they enter the safety exit. After each trial, the subject is asked to report how difficult they found it to escape the virtual predator (4s). B) Modified schematic representation from the model proposed by Ydenberg and Dill [1]. As the distance between the prey and the predator decreases, the cost of fleeing decays, while the cost of not fleeing rises. D* represents an optimal point where the prey should flee. C) Histograms showing the distribution of subjects’ FID choices for early, mid and late attacking predators respectively. The X axis represents FID, while the Y axis represents frequency of choice.

## Results

### Behavior

We first examined the behavioral data by applying a repeated-measure three-way ANOVA (Predator Type X Reward Level X Shock Level) for escape responses (e.g. FIDs). Results showed a main effect of predator type (F(2,54) = 82.59, p < .001). Posthoc comparisons for the predator type X shock level interaction revealed that the difference in FID choices between high and low shock levels only exist in the slow attacking predator condition (p = 0.013). This shows that subjects considered the level of potential danger while choosing FID (more risk-averse when shock is higher), but only when considering the slow attacking threat where there was time for strategic planning. The same repeated-measure three-way ANOVA was performed for escape difficulty ratings. A main effect of predator type was found (F(2,54) = 49.77, p < .001), showing that subjects estimate fast attacking predator as the most difficult predator type to escape (all posthoc comparisons: p < .001). Significant interactions were found for predator type X shock level (F(2,54) = 13.68, p < .001) and predator Type X reward level (F(2,54) = 4.39, p = .017). For the predator type X shock level interaction, we found that rating was higher in the high shock condition, but only in the slow attacking predator (posthoc comparison: p < .001). This is intriguing because the predator’s attack distance is identical in both shock levels, yet subjects perceived the threat to be more difficult to escape in the high shock condition (see Fig. S1).

### Neural basis of fast and slow escape decisions

We next investigated the neural basis of escape decisions for the fast and slow attacking threat. In order to control for timing differences between conditions, besides modelling the rest of the trial as a boxcar function, we specifically looked at the 2s before the FID bottom press as a period where subjects form their final decisions. We chose to time lock 2s before the flight decision for two main reasons: (i) because it allowed us to examine the neural ramping up of the flight decision, and controlled for the contamination of outcome; (ii) reduced the amount or trials that would be lost for the fast attack condition. Also, to control for any confounds of pain, we excluded the caught trials (Number of caught trials: far attacking predator: mean = 8 ± SD =3; mid attacking predator: mean =5 ± 2; near attacking predator: mean = 4 ±1), using these events as regressors of no interest. As the mid AD condition was a priori used an anchor for the fast and slow AD threats, we focus on activity for the fast and slow AD (see SOM for further results). A whole brain analysis was first performed to locate regions associated with decisions under reactive fear (fast attacking predators) and cognitive fear (slow attacking predators). Detailed regions of activation can be found in Table S1 – S2. As shown in Fig. 2, with data extracted from independent anatomical ROIs differential activation patterns were clearly found for the different predator ADs (See SOM for a full list of activated regions). In order to confirm the dissociation between the reactive and cognitive fear systems (represented by PAG and vmPFC respectively), we computed a two-way ANOVA (region X predator type) using signal change drawn from independent ROIs from PAG and vmPFC. There was a main effect of region (F = 5.77, p = .017) and a significant interaction between region and predator type (F = 11.50, p < .001). For the Fast AD > Control contrast, we observed increased activity in PAG and MCC. A direct comparison between high vs low shock for the fast AD predator revealed increased activity in the PAG, suggesting that PAG is evoked when the threat is high (see table S7).

**Fig. 2.**
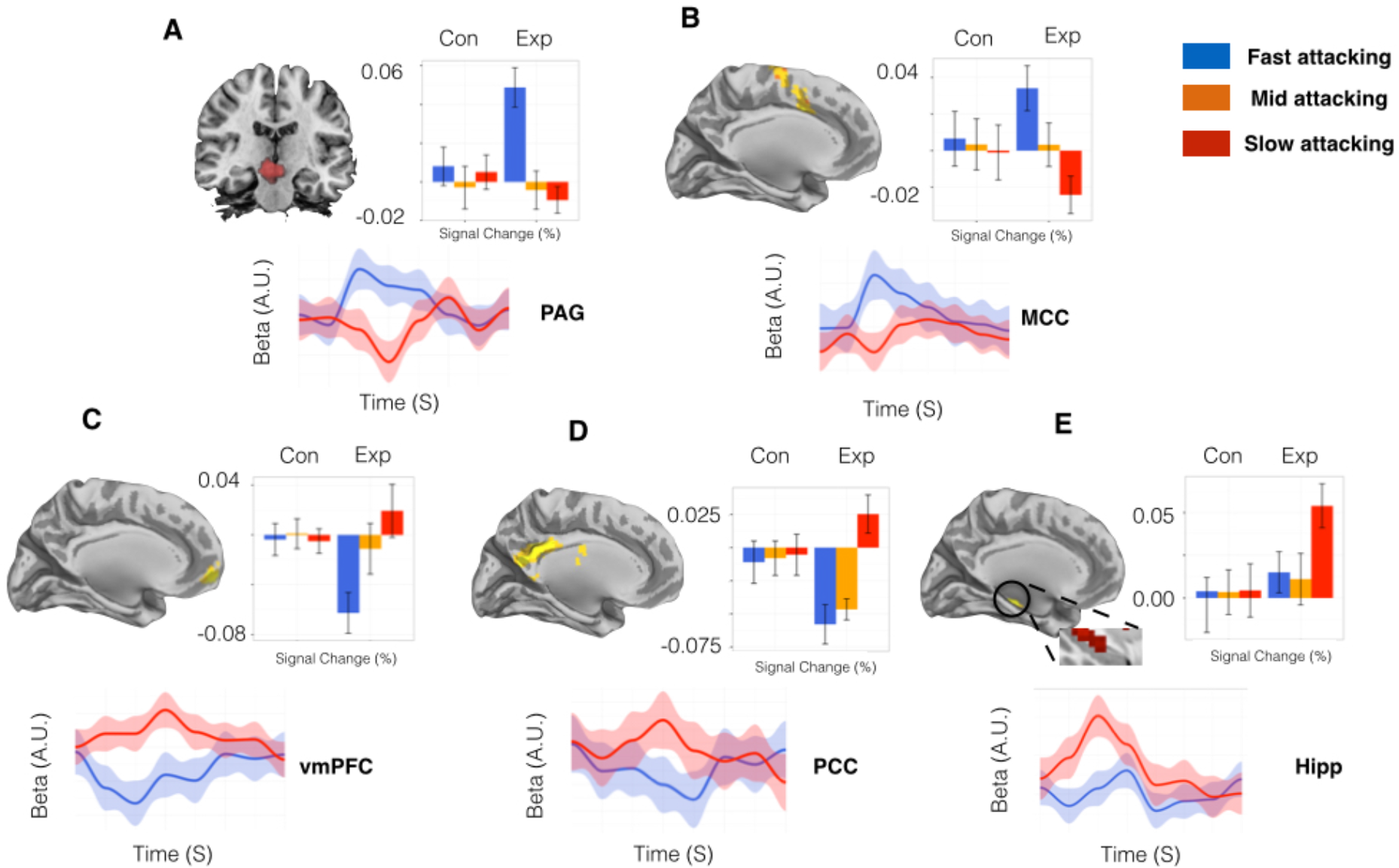
Main regions of interest and signal changes associated with fast and slower attacking threats. Parameter estimates and time series extracted from A) midbrain, B) MCC, C) vmPFC D) PCC and E) the hippocampus. Activations shown in the graph show clusters from the whole brain activation, while the signal change data were extracted from independent anatomical ROIs. The upper graph displays parameter estimates. Y axis represents percent signal changes, and X axis is the predator type. The lower graph display time series extracted in a course of 16 seconds. Blue line: Fast predator; Red line: Slow predator. Beginning of the time series represents the time point when the FID event comes online.

On the other hand, the slow AD > Control revealed increased activity in the cognitive fear circuitry including the vmPFC, PCC and the hippocampus. While no amygdala was observed for the main contrast, we did find that a direct comparison between high vs low shock for the slow AD predator, showed increased activity in the amygdala and hippocampus (see table S8). To further disentangle the effect and increase the sensitivity of the analysis, we extracted the signal changes and BOLD signal time series from the predefined ROIS (i.e. PAG, MCC, vmPFC, PCC, hippocampus), regions that have previously been associated with fear, anxiety, and decision making under stress (Mobbs et al., 2007). A conjunction between fast and slow attacking threats showed that the medial dorsal thalamus (MDT) was commonly activated. Although this is an exploratory finding, it is intriguingly because MDT is directly or indirectly connected to both fear circuits, stimulation of the MDT results in depression or potentiation of both circuits and it is thought to play a role in behavioral flexibility (Krout and Loewy, 2000; Vertes et al., 2015).

### Computations that support escape decisions

To explore how the observed FIDs might be understood in terms of rational decision making about the costs and benefits of flight, we developed a simple Bayesian decision theoretic model of the task. The process by which subjects make escape choices under different predator ADs can be decomposed to two steps: 1) Predicting the different predators’ distribution of attack distances, by learning from experience and 2) Choosing a FID by comparing the money obtained against the risk of shock for each possible FID, in expectation over the predicted attack distance distribution and informed by the individual’s subjective preference (utility) levels for shock vs. money. For simplicity, we assume a Bayesian ideal observer model of subjects’ learning to estimate the attack distances of different predators from trial-by-trial experience, and that FID choices are then determined (with softmax noise) by computing the expected utility (money minus shock) for each possible escape distance. Details of the model can be found in the online supporting material. We then calculated the distance between utility resulted from subjects’ actual FID and the predicted Bayesian ideal FID, which is a measure of optimal performance (Fig. 3).

**Fig. 3.**
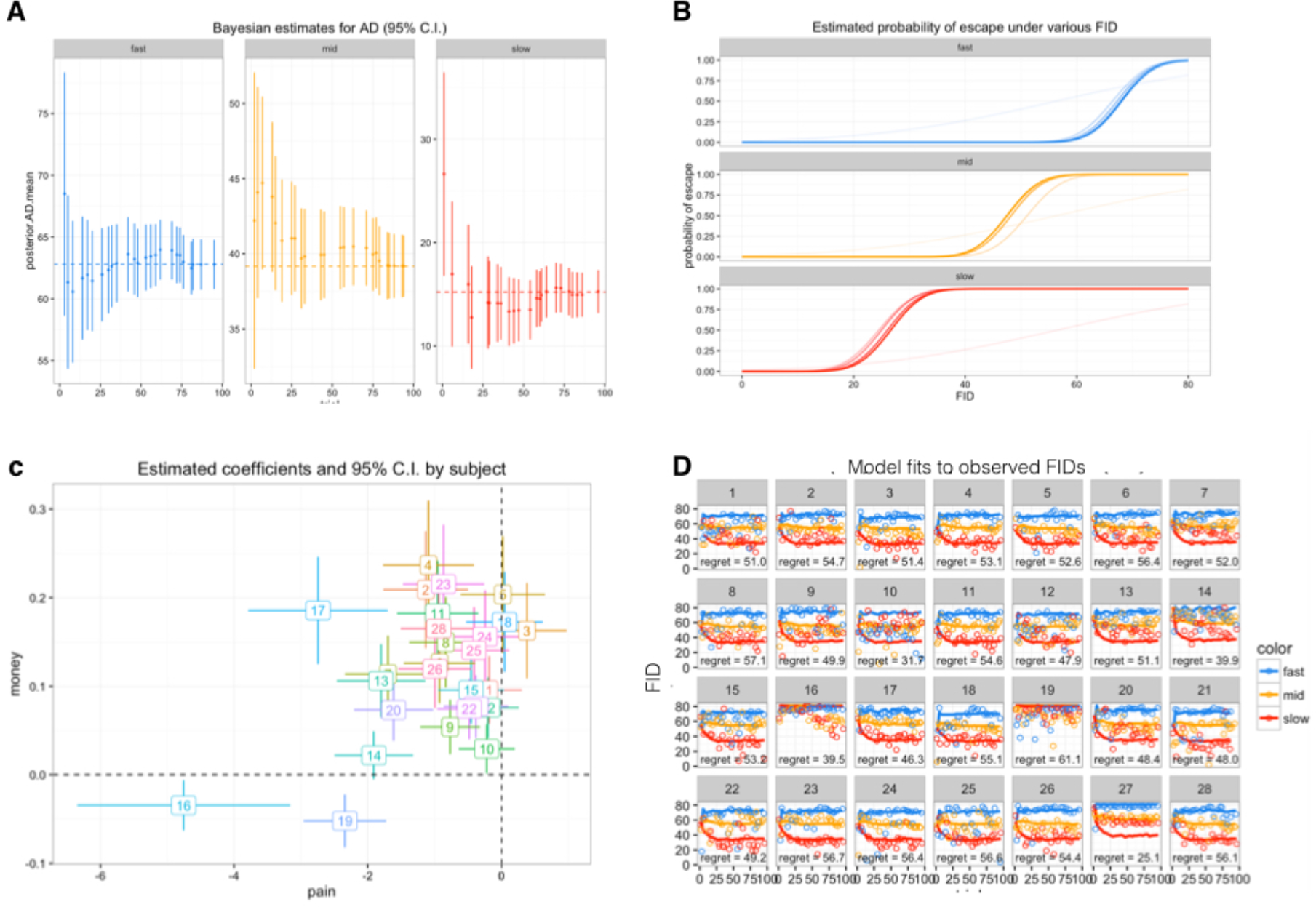
Visualization of Bayesian Modelling Results. (A) Bayesian ideal observer estimates of predator AD, based on the unknown-mean-known-variance ideal learner model, as a function of experience in the task. Color 1 (blue), 2 (yellow), 3 (red), 4 (green) corresponds to fast, mid, slow and control predators, respectively. The graph shows the observer’s 95% credible interval almost always contains the true mean, indicating an appropriate modeling of uncertainty. (B) Relationship between FID and the chance of escape for each predator type. Chance of escape increases with FID, but with different growth patterns in every predator type. (C) Estimated coefficients for each subject, along with 95% confidence intervals. X axis represents the pain coefficient β_1 in the utility function, and Y axis represents the monetary reward coefficient β_2. For a rational player, β_2 should be positive (seeking money) and β_1 should be negative (avoiding shock). (D) Model fits to observed FIDs. X axis represents trial numbers, and Y axis represents FID. Ideal FID choices predicted by the ideal Bayesian observer (lines), subjects’ actual FID choice (dots), and the difference between them in terms of utility under estimated individual preference. Regrets here represents the utility difference between actual chosen FID and the ideal Bayesian choice.

We next examined neural circuits correlated with subjects’ preference parameters in the Bayesian decision making model. For a rational player, the preference for reward should be positive, while the preference for shock should be negative. Thus, greater reward or shock sensitivity here corresponds to larger (positive) *β*_2_ and smaller (more negative) *β*_1_. The parametric modulation analysis over the [predator > control] contrasts revealed that, for the fast predator condition, higher reward sensitivity was associated with activations in bilateral putamen, while higher shock sensitivity is associated with engagement in PAG and bilateral insula. On the other hand, for the slow predator condition, right caudate was found to be associated with higher reward sensitivity, while PCC was found to be associated with higher shock sensitivity. A display of the activated regions can be found in Fig. S3. A detailed layout of the activated regions can be found in table S9 and table S10.

Next, in order to investigate what neural circuits are responsible for the optimization of escape decision making, we considered a measure of performance optimality related to the per-trial spread between subjects’ actual and Bayesian ideal FIDs. In particular, we computed the difference between the actual trial-specific utility *U*(*FID*) and the maximum (Bayesian optimal) utility the subject could possibly get on the trial (*U*(*FID*)_*max*_), given their estimated subjective utilities. A smaller difference (e.g., less regret relative to ideal) implies more consistent Bayesian decision making; variation around the ideal FID will increase the difference. The difference on every trial were entered as a subject-level parametric modulator separately under each [Predator > Control] conditions. For the far predator condition, we found that better Bayesian decision making (smaller distance to ideal) was associated with activity in MCC, middle frontal gyrus, and superior motor cortex. On the other hand, better Bayesian decision making in the slow AD condition was found to be associated with activity in bilateral hippocampus, as shown in Fig. 4.

**Fig. 4.**
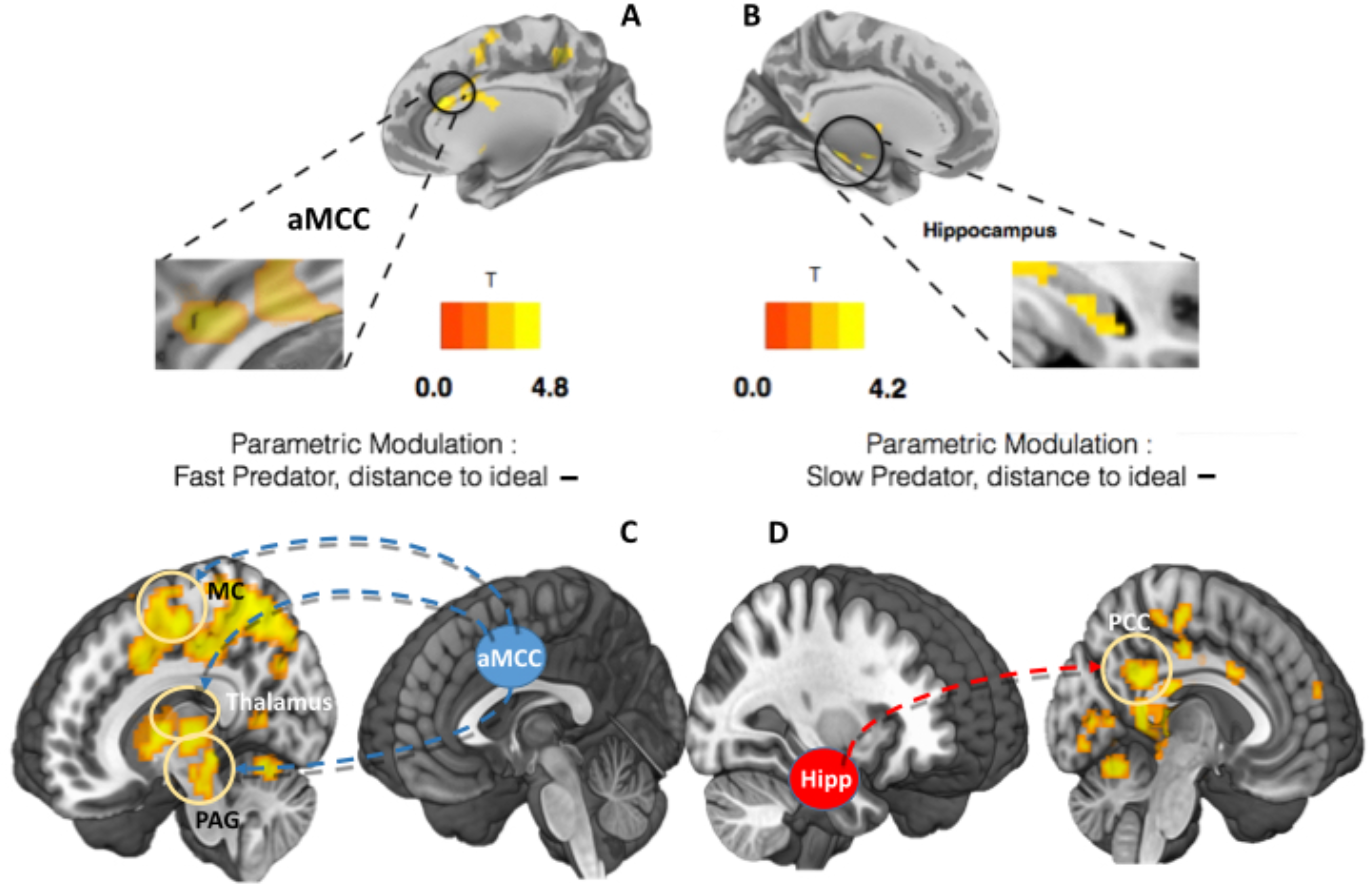
Regions, and their connectivity, associated with parametric modulation of “distance to ideal”. (A) Brain regions associated with increased Bayesian decision optimality in the fast AD condition. Better decision making was associated with increased activity in MCC and superior motor cortex. (B) Brain regions associated with decreased distance (increased Bayesian decision optimality) in the slow AD condition activated regions include bilateral hippocampus and bilateral caudate. A display of the correlation results can be found in Fig. S4. (C) Connectivity analysis using MCC as seed over the contrast [fast predator > control]. Positive connectivity was found between MCC, motor cortex (MC), thalamus and the PAG. (D) Using the hippocampus as seed over the contrast (slow predator > control), positive connectivity was found between the hippocampus and PCC.

### Functional connectivity between computationally defined regions

To investigate the interplay among the brain regions involved in escape decision optimization, a functional connectivity analysis was performed for the response phase (escape decision) using a generalized psychophysiological interactions (PPI) approach (McLaren et al., 2012); In order to confirm the patterns observed in the whole brain flexible model, we first adopted independent seed regions of MCC and hippocampus from previous research (Mobbs et al., 2007). For the contrast of Fast Predator > Control, we showed a significant coupling between the MCC seed, the PAG, motor cortex and bilateral thalamus. For the contrast of slow Predator > Control, we showed a significant coupling between the hippocampus seed and PCC. This suggest that when the subjects are provided time for decision flexibility, they use a search and employ approach which prepares them for action, as shown in figure 4.

## Discussion

We have demonstrated that subjects apply different nodes of the survival circuitry when escaping fast or slow attacking threats. Our analysis revealed increased activity in ‘reactive fear’ circuits namely the PAG and the MCC for the fast AD predator, regions that are implicated in response to fast and imminent threats. Supporting comparative work (An et al., 1998), connectivity analysis revealed a significant couple between the MCC and PAG and recent animal work has shown that the optogenetic activation of glutamatergic neurons in the dorsal lateral PAG induce motor responses (e.g. flight; Tovote et al., 2016). The MCC is also a critical component of the defensive survival circuitry and has afferent projections to the ventral striatum, receives efferents from the medial dorsal thalamus and has bidirectional projections with the amygdala (Shackman et al., 2011). It has also been suggested that control signals in the MCC may resolve conflict between defensive strategies (e.g. freezing or fleeing). This has led to the theory that the cells in the MCC are involved in linking motor centers with defensive circuits (Shackman et al., 2011).

Our analysis for the slow attacking threat contrast revealed activation in three key areas of the ‘cognitive fear’ circuitry: the vmPFC, hippocampus and PCC. Structural and function connectivity between these structures has been shown in humans and primates, supporting conserved pathways across species (Price, 2005). Primate research has found that the primate PCC responds to risky decision-making and scales with the degree of risk (McCoy and Platt, 2005). The PCC is also correlated with a salience signal reflecting the deviation from the standard option, suggesting a role in the flexible allocation of neural resources (Heilbronner et al., 2011). The goal of the PCC may be to harvest information for escape decisions under conditions of protracted threat. This fits with the proposal that through its connections with the hippocampus, the PCC may integrate memory guided decisions with current decision processes which may involve a ‘preparation for action’ through rumination and worry.

The vmPFC is also a key player in the defensive survival circuitry. Single-cell recordings in rodents have shown that the mPFC contains ‘strategy selective’ cells, which are thought to be involved in the coordination of defensive responses (Halladay and Blair, 2015). This fits with the idea that the mPFC plays a role in selecting adaptive strategies that are mapped onto motor responses. Indeed, work in humans shows that larger buffer distances are associated with activity in the vmPFC and decreased activity in this regions is associated with panic related motor actions (Mobbs et al., 2010; Perkins et al., 2015). Our data builds on these findings by showing that vmPFC, hippocampus and PCC form a strategic and flexible decisions when the agent has time to contemplate the best escape action. Our findings support the role of the survival circuitry by proposing that ‘fast’ fear reactions engage a neural circuits involved in reactive fear, while ‘slower’ escape decisions are associated with strategic avoidance and conscious cognitive fear (LeDoux and Pine, 2016).

Our Bayesian model also provides insights into how the distinct regions of the survival circuits associated with optimal escape. Two core regions where associated with optimal escape. The MCC for the fast attacking threat and the hippocampus for the slow attacking threat. While it is accepted that the PAG needs input to make optimal decisions, it is unclear where this input comes from. A few candidates exist, among them is the MCC. The MCC is highly connected to the lateral PAG and according to adaptive control theory is a “central hub” where information about reinforcements are passed to motor control areas to coordinate goal directed behaviors (Shackman et al., 2011). Our connectivity results support this conclusion showing that the MCC was coupled with activity in the PAG and the motor cortex. This proposes that the MCC is one candidate region for the integration of current goals and implement aversively motivated instrumental motor behaviors (i.e. when to flee a threat (Shackman et al., 2011)).

The hippocampus has been identified as a central processing regions involved in approach-avoidance conflict and corresponding choice monitoring (Loh et al., 2016). Theorists have proposed that the hippocampus computes comparators that assess multiple goals and in turn correct actions (McNaughton and Gray, 2000) possibly through a flexible constructive process involved in problem solving (Hassabis and Maguire, 2007). When there is time to gather information, the hippocampus may play a role in drawing on previous threat encounters to optimize current actions. Indeed, the hippocampus plays a role in spatial and temporal ‘where’ and ‘when’ memory and has theoretically been linked to escape decisions (Litvin et al., 2015) and may possibly by resolving conflict between fitness promoting behaviors (e.g. escape vs reward;(Bach et al., 2014). The ventral hippocampus is particularly interesting given its role in emotion. The ventral hippocampus is connected to the vmPFC, basolateral amygdala and the bed nucleus of the stria terminalis and is believed to contribute to the control of behavior under anxiogenic conditions (Bannerman et al., 2004). Our connectivity results support this conclusion showing that the hippocampus was also coupled with activity in the PCC, a region thought to be involved in adaptive decisions (Pearson et al., 2011). Together, these theories support a Bayesian role for the hippocampus, where it gathers information to optimize directed escape during slow, but not fast, attacking threat.

In summary, we introduce a paradigm that allows researchers to map escape decisions onto the defensive survival circuitry. This circuit can be separated into a fast ‘reactive fear’ circuit involved in escape decision when time is limited and ‘slower ‘cognitive fear’ circuits are involved in the strategic avoidance and conscious feelings of fear when the agent has time to think. More specifically, sections of these circuits differentially optimize escape decisions with the MCC centered on making fast decisions associated with imminent threat and the hippocampus in slow strategic decisions that are characterized by protracted threat assessment. These results provide a new window in the role of the defensive survival circuitry in adaptive escape decisions and transform the way we view the neural circuits involved in human fear.

## Methods

### Participants

A total number of 30 subjects completed informed consent in accordance with the guide lines of the Columbia University Institutional Review Board and were remunerated for their participation. Data from 1 subject was lost due to computer error. One additional subject was excluded due to excessive movement during the scan. Our final sample consisted of 28 subjects (17 women, Age = 25.4 ± 7.3 years).

### Experiment Design

Subjects were scanned while they viewed stimuli on the screen. The screen displayed a 2D run way, with a virtual predator “attacking” from the left entrance. In the current paradigm, the goal of the subject was to escape the attack from a certain virtual predator, by pressing a bottom at the desired timing. Once the bottom was pressed, a triangle representing the subject started moving towards a “safety exit”. Subjects gained reward if they escaped to the safety before the predator caught them; On the other hand, they were given a mildly aversive electric shock if they were caught. The goal of the game was to choose the right time to flee meaning acquiring the maximum amount of reward while still escaping the virtual predator. To accomplish this, reward in each trial scaled with time spent before pressing the bottom. The longer subjects stay (the smaller the FID), the more reward they get. But staying too long could mean receiving an electric shock from the predator.

The runway has a total length of 90 units, where the prey is placed 10 unites to the safety exit. While in the approaching mode, the predator proceeds with a speed of 4 units/second; while in the chasing mode, the predator proceeds with a speed of 10 unites/second. The reward participants get on every trial is positively correlated with the length they stay. A total number of 96 trials exist in the current paradigm, factorial divided to cover different predator attack distances, shock level and reward levels (3 X 2 X 2). Subjects were first presented with a screen indicating which type of predator and shock/reward level will be presented in the next trial for 2s. This shock/reward indicator informs the next 4 trials. Next the trial begins, where subjects observes an artificial predator slowly looms towards their triangle. After a designated time period, which is learned by the participant, the artificial predator will attack by speeding up (i.e. the attack distance) when it reaches a designated position. There are 4 types of attacking predators, with their attack distances subject to 3 Gaussian distributions (i.e. fast attacking predator, mid attacking predator, slow attacking predator, and control predator; See Fig.1 for details). As the threat slowly looms towards them, the subjects determine their initiation of flight by pressing the FID button. To make sure reaction time plays no role in FIDs, we manipulated the speed of the predator and the subject so that once the threat speeds up to attack, it is impossible to escape. After the trial, subjects are required to rate the difficulty of escape using a visual analogue (1-5) scale. A detailed breakdown of the experimental procedure can be seen in Fig.1. The experiment consists of a total number of 96 trials. After 48 trials, the assignment of predator-color relationship was altered to introduce novelty and avoid the fixation of subject strategies.

Before the start of the actual experiment, subjects went through a brief practice session of 8 trials to familiarize themselves with the paradigm. In the practice session, subjects played the same game, but the predators’ attack distances were drawn from different distributions than the ones in the actual experiment.

### Behavioral Analysis

Due to the relative simplicity of our task and exposure to a practice session, subjects’ performance reached saturation very quickly after the beginning of the experiment. By “saturation”, subjects quickly formed their own patterns of choice making and carry through the rest of the experiment. Thus, instead of looking at trial-by-trial changes of the FID, we focus on the differences of FID between different predator conditions, and approaches subjects’ learning behavior by Bayesian modeling (described later).

Subjects’ choice of FID, reward from the trial, and escapability ratings were collected on each trial. We used repeated-measures three-way ANOVAs (of Predator Type X Reward Level X Shock Level) to assess differences in FID, reward and escapability ratings between the various conditions.

### Acquisition and Analysis of fMRI data

All fMRI data were acquired using a GE Discovery MR750 3.0 T scanner with 32-channel headcoil. The imaging session consisted of two function scans, each twenty minutes, as well as a high-resolution anatomical Ti-weighted image (1mm isotropic resolution) collected at the beginning of each scan session. For functional imaging, interleaved T2*-weighted gradient-echo echo planar imaging (EPI) sequences were used to produce 45 3-mm-thick oblique axial slices (TR = 2 sec., TE = 25 ms, flip angle = 77°, FOV = 192 x 192 mm, matrix = 64 x 64). Each functional run began with five volumes (1000 msec) before the first stimulus onset. These volumes were discarded before entering analysis to allow for magnetic field equilibration. Stimulus were presented using Cogent (matlab-based package). Participants viewed the screen via a mirror mounted on the head coil, and a pillow and foam cushions were placed inside the coil to minimize head movement.

Analysis of fMRI data was carried out using scripted batches in SPM8 software (Welcome Trust Centre for Neuroimaging, Lon-don, UK; http://www/fil.ion.ucl.ac.uk/spm) implemented in Matlab 7 (The MathWorks Inc., Natick MA). Structural images were subjected to the unified segmentation algorithm implemented in SPM8, yielding discrete cosine transform spatial warping coefficients used to normalize each individual’s data into MNI space. Functional data were first corrected for slice timing difference, and subsequently realigned to account for head movements. Normalized data were finally smoothed with a 6-mm FWHM Gaussian kernel.

Preprocessed imaged were subjected to a two-level general linear model using SPM8. The first level contained the following regressors of interest, each convolved with the canonical two-gamma hemodynamic response function: a 2-second box-car function for the onset of the trial (where the color of the incoming predator is shown); a 4-8 second (duration jittered) box-car function from the onset to 2s before when subjects make the flight decision; a 2-second box-car function for the phase before subjects make the flight decision; a 4-8 second (duration jittered) box-car function for the remainder of the trial. Mean-centered trait anxiety ratings, escapability ratings and parameters in the Bayesian decision model were included as orthogonal regressors. In addition, regressors of no interest consisted of motion parameters determined during preprocessing, their first temporal derivative and discrete cosine transform-based temporal low frequency drift regressors with a cutoff of 192-seconds.

Beta maps were used to create linear contrast maps, which were then subjected to second-level, random-effects one-sample t tests. In Addition, A flexible factorial model was used to examine the main effects of predator type, reward level and shock level. Interaction effects between predator type, reward level and shock level were also examined using the factorial model. The resulting statistical maps were thresholded at P < 0.05 corrected for multiple comparisons (false discovery rate [FDR] corrected (Genovese et al., 2002)). A flexible factorial model was used to examine the interaction effects between predator type, reward level and shock level. The threshold for those specific contrasts was set at p < 0.05 (FDR corrected).

A hypothesis driven region of interest (ROI) analysis were performed after the whole brain analysis for regions with strong a priori spatial hypotheses. The ROI analysis was performed using regions associated with the processing of fear, threat and decision making. Independent ROIs were chosen from previous research showing similar effects(Mobbs et al., 2007; Tedeschi et al., 2015). The threshold for these analyses was set at p < 0.05, small volume correction (SVC).

The functional connectivity analysis was performed for the response phase (escape decision) using a generalized psychophysiological interactions (PPI) approach. The connectivity analysis was carried out based on the [predator condition > control condition] contrast.

### Bayesian Decision Making Model

In order to better understand and characterize the process by which subjects learn the nature of the predators and the subsequent adjustment of their strategies, we applied a Bayesian learning model to simulate how subjects adapt to attacks from different predators. In the FID task, each predator type (far, mid and close) had a predefined location of attack. These attack distances were subject to three Gaussian distributions, corresponding to the three predator types, as mentioned before. As we shall see in the following section, the modeled *ideal Bayesian learner* will update its belief (posterior) about the next attacking distance by combining observations (likelihood) of attacks and an earlier belief (prior) using the Bayes rule, which is the statistical principle of reasoning with evidence and belief. The updated belief will then inform the optimal choice of the flight initiation distance (FID), in order to win money and avoid shock.

Naturally, subjects base their flight choices on the *predicted* attacking distance of the current trial. At first, subjects have zero to minimum knowledge about the color-predator associations, and each predator’s defined range of attacks. However, as the experiment progresses, subjects accumulate knowledge about the predators, and form better predictions of the attack distance for the current trial after seeing the corresponding predator color. Thus, the task is modeled as a process where subjects learn towards the true distribution of the attack distance.

A priori to observing attacking, the attack distance of a certain predator is believed to be drawn from a Gaussian distribution *AD*|*c* ~ *N*(*µ*^*c*^,*σ*^2^), where *c* represents the predator type. At the start of the experiment, the mean parameters are unknown and hence assumed to follow the same prior distribution. Here we adopt the conjugate prior distribution 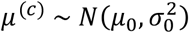 with a large variance to reflect minimum prior knowledge. Meanwhile, we assume the variance of likelihood (*σ*^2^) to be known, because in the practice phase subjects have already been exposed to predators with identical AD variance as in the formal experiment.

Upon observing attacks, the posterior distribution for the mean parameter is updated by the Bayes rule, yielding:

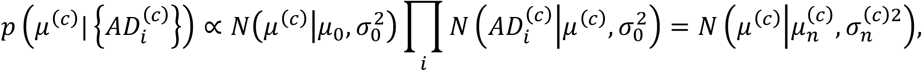

where 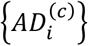 are the *observed* distances of a total number of *n* attacks from type-*c* predators. The posterior is also a Gaussian, with parameters updated through 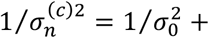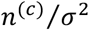 and 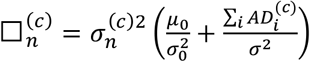. This posterior of the mean parameter directly *induces* the (posterior) predictive distribution of the upcoming attack distance, given by

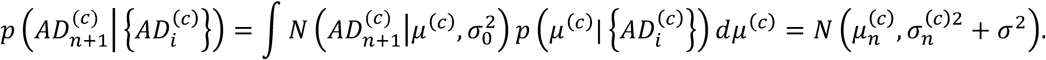

An ideal Bayesian learner will base its FID choice on this distribution. Now, under the context of the current paradigm, a subject chooses FID from a finite set of options by trading off two critical factors: the *risk of getting shocked* and the *monetary reward*. With a large FID, risk is reduced while less reward will be given; with a small FID, the opposite. We then define an overall utility as a weighted combination of the two factors:

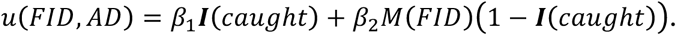

The coefficients *β*_1_, *β*_2_ are individuals-specific weights to adjust the preference between the two factors. ***I***(*caught*) is the indicator function for the event of getting caught and killed (evaluates to 1 if caught, 0 otherwise); and *M*(*FID*) is the amount of money rewarded if escape is successful. This utility is a random function since getting caught is a random event --- the optimal decision should then be based on the *expected value* of utility, namely the *Bayesian risk*, which is estimated from the latest posterior predictive distribution and takes the form of

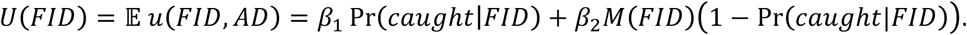

Here the probability of being caught Pr(*caught*|*FID*) can be computed by solving a simple chasing problem, where the actual speed of the particular predator and subject were taken into computation to determine if the subject would be caught or not.

The Bayesian learner’s optimal choice is set as a reference to measure the performance of subjects. Clearly, the optimal FID is one that maximizes *U*(*FID*). Yet, the behavior of a human player is also influenced by unobserved factors such as personality traits and is not necessarily Bayesian optimal. To quantify individual differences of decision making through coefficients (*β*_1_, *β*_2_), we fit a discrete choice model (multinomial logit) for each subject, which assumes that the probability of picking an option is proportional to the corresponding exponential utility:

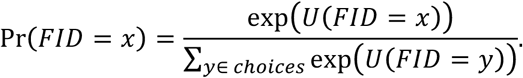

The coefficients can be estimated by maximizing the overall likelihood, namely

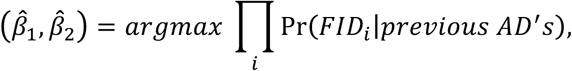

where the product is taken over trials. By fitting to our data, a subject can be quantified by the corresponding coefficients:

*β*_1_ (pain): expected to be non-positive, and a bigger absolute value means stronger aversion towards risk and its associated penalty (electric shock).
*β*_2_ (money): expected to be non-negative, and a bigger value measures stronger favor of monetary reward.
−*β*_1_/*β*_2_: the relative weight between the two factors, which conceptually means the maximum amount of money one is willing to pay to avoid being caught (and shocked) once

Those parameters are then entered into fMRI parametric modulation analysis to determine the brain regions where signals covariate with the parameters. Results of the modulation analysis are discussed in the next section.

On every particular trial, the difference between the actual utility *U*(*FID*) and the Bayesian optimal utility *U*(*FID*)_*max*_ is calculated as a measure of choice optimality (Alternatively, the “utility” used to calculated differences here can be “normalized” by dividing out *β*_2_, the reward preference parameter.

## Acknowledgments

The authors wish to thank John O’Doherty, Peter Dayan, Alex Shackman, James Curley, Joe LeDoux for their advice and Nir Jacoby for his work on an earlier version of the paradigm. This study was supported by a grant from the NARSAD to DM.

## Author Contributions

DM, SQ, JS, DH, and ND designed the study. SQ collected the data. DM, SQ, ND, FG, analyzed the data. DM, SQ, ND, FG, DH wrote the paper.

---

**Fig. S1.**
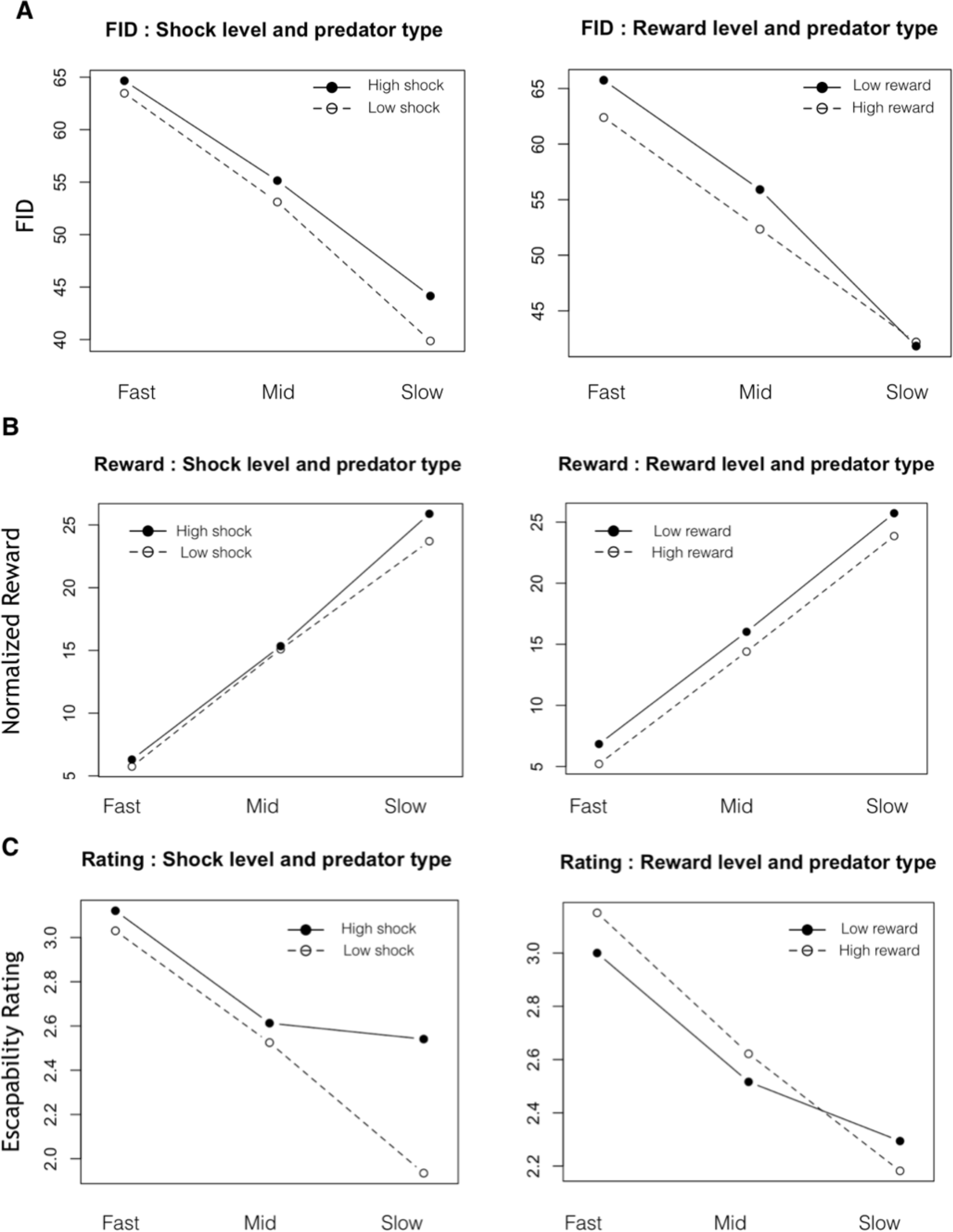
ANOVA for A) FID, B) Reward and C) Escapability rating.

**Fig. S2.**
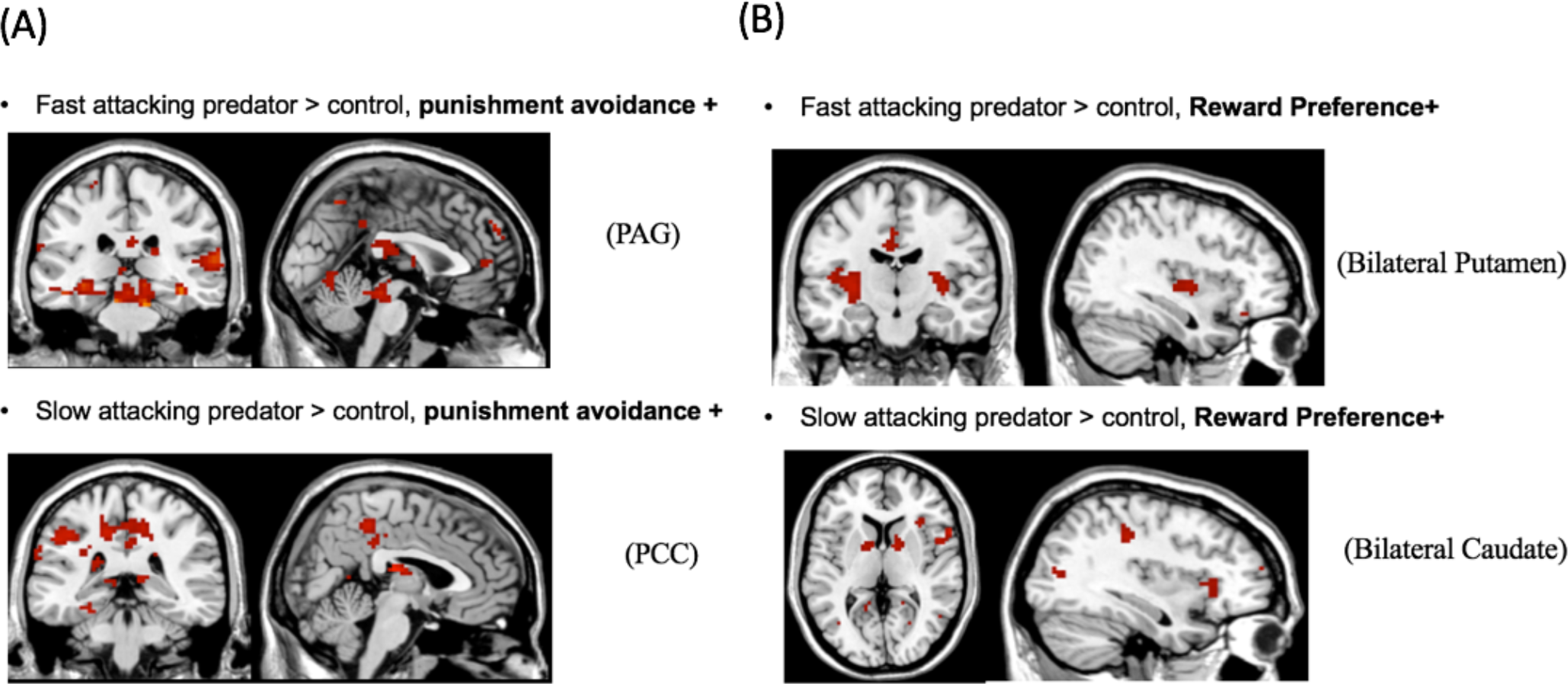
Activated regions for 1st level parametric modulation with reward and punishment sensitivity parameters in the Bayesian decision making model.

**Table S1.** Activation Table for Contrast [Far Predator > Control] (Whole Brain)

**\* P<0.05, FDR corrected**
| Brain Region | Left/Right | Cluster Size | t-score | Coordinates |  |  |
| --- | --- | --- | --- | --- | --- | --- |
|  |  |  |  | x | y | z |
| *Far Predator > Control* |  |  |  |  |  |  |
| Middle temporal Gyrus | R | 461 | 7.82 | 48 | -66 | 6 |
| Middle temporal Gyrus | L | 155 | 6.19 | -42 | -69 | 6 |
| Precuneus | R | 302 | 5.99 | 6 | -48 | 48 |
| Precentral Gyrus | L | 81 | 4.52 | -42 | -9 | 48 |
| Superior Temporal Gyrus | L | 94 | 5.11 | -57 | -42 | 21 |
| Insula | L | 199 | 4.96 | -33 | 21 | 3 |
| Insula | R | 285 | 6.72 | 37 | 20 | -1 |
| Mid Cingulate Gyrus | L | 71 | 4.36 | -12 | -21 | 39 |
| Midbrain |  | 244 | 6.51 | 3 | -28 | -12 |
| Supplementary Motor Area | R | 357 | 6.28 | 17 | 5 | 60 |
| Caudate | R | 29 | 4.67 | 8 | 10 | -5 |

**Table S2.**
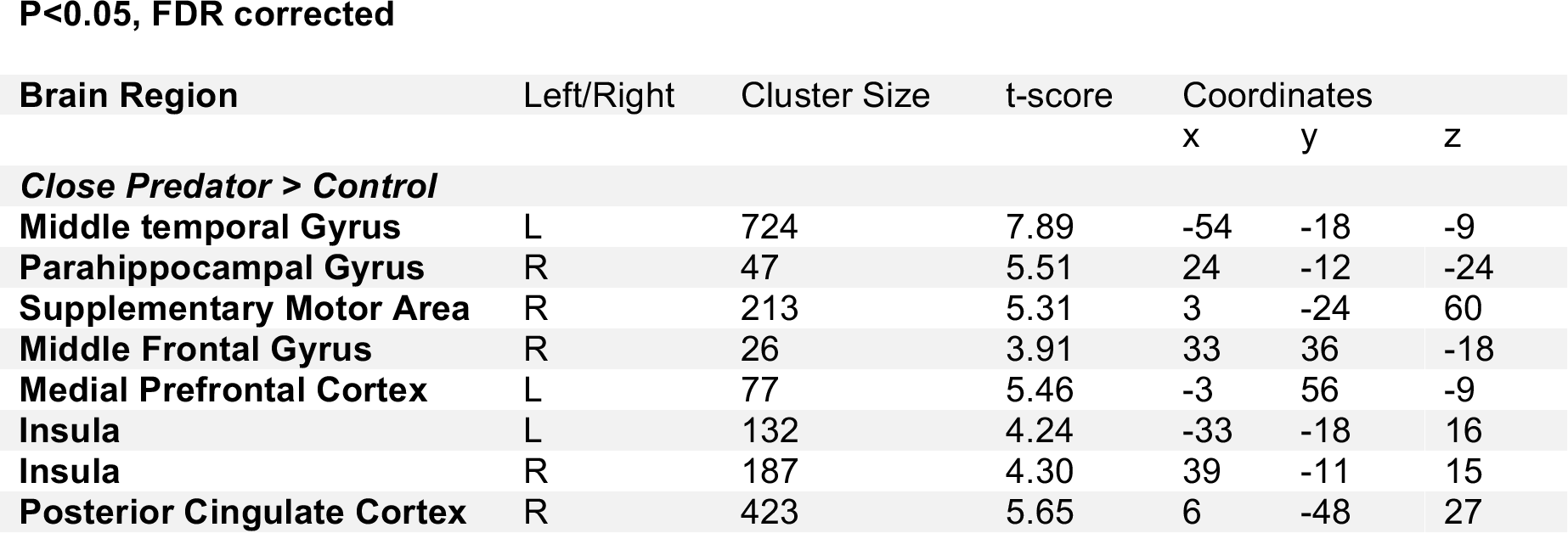
Activation Table for Contrast [Close Predator > Control] (Whole Brain)

**P<0.05, FDR corrected**
| Brain Region | Left/Right | Cluster Size | t-score | Coordinates |  |  |
| --- | --- | --- | --- | --- | --- | --- |
|  |  |  |  | x | y | z |
| *Close Predator > Control* |  |  |  |  |  |  |
| Middle temporal Gyrus | L | 724 | 7.89 | -54 | -18 | -9 |
| Parahippocampal Gyrus | R | 47 | 5.51 | 24 | -12 | -24 |
| Supplementary Motor Area | R | 213 | 5.31 | 3 | -24 | 60 |
| Middle Frontal Gyrus | R | 26 | 3.91 | 33 | 36 | -18 |
| Medial Prefrontal Cortex | L | 77 | 5.46 | -3 | 56 | -9 |
| Insula | L | 132 | 4.24 | -33 | -18 | 16 |
| Insula | R | 187 | 4.30 | 39 | -11 | 15 |
| Posterior Cingulate Cortex | R | 423 | 5.65 | 6 | -48 | 27 |

**Table S3.** Activation Table for Trial-bytrial parametric analysis with FID.

**P<0.05, FDR corrected**
| Brain Region | Left/Right | Cluster Size | t-score | Coordinates |  |  |
| --- | --- | --- | --- | --- | --- | --- |
|  |  |  |  | x | y | z |
| ***FID parametric modulation + : Far Predator*** |  |  |  |  |  |  |
| Midbrain | L | 280 | 7.80 | -12 | 27 | -3 |
| Hippocampus | R | 31 | 7.03 | 18 | -13 | -13 |
| Thalamus | L | 90 | 6.87 | -11 | -26 | -1 |
| Insula | L | 63 | 5.23 | -38 | 5 | -12 |
| MCC | L | 105 | 6.94 | -3 | 2 | 42 |
| ***FID parametric modulation - : Far Predator*** |  |  |  |  |  |  |
| Middle Temporal Gyrus | L | 51 | 3.63 | -51 | -15 | -15 |
| Middle Temporal Gyrus | R | 40 | 3.45 | 51 | -27 | -15 |
| Middle Occipital Gyrus | R | 47 | 4.07 | 30 | -90 | 15 |
| ***FID parametric modulation + : Close Predator*** |  |  |  |  |  |  |
| Inferior Frontal Gyrus | R | 64 | 4.79 | 36 | 21 | -6 |
| Middle Frontal Gyrus | R | 93 | 5.76 | 39 | 60 | 3 |
| MCC | L | 79 | 4.73 | -1 | 33 | 36 |
| Cerebrum | R | 66 | 7.88 | 24 | -9 | 36 |
| ***FID parametric modulation - : Close Predator*** |  |  |  |  |  |  |
| Superior Temporal Gyrus | L | 29 | 5.56 | -57 | -51 | 12 |
| Inferior Frontal Gyrus | L | 43 | 6.46 | -54 | 27 | 18 |
| PCC | L | 33 | 5.19 | -9 | -51 | 27 |

**Table S4.** Activation Table for Trial-by-trial parametric analysis with Escapability.

**P<0.05, FDR corrected**
| Brain Region | Left/Right | Cluster Size | t-score | Coordinates |  |  |
| --- | --- | --- | --- | --- | --- | --- |
|  |  |  |  | x | y | z |
| Escapability parametric modulation + : Fast Predator |  |  |  |  |  |  |
| Middle Temporal Gyrus | L | 104 | 7.52 | 42 | -51 | -21 |
| Midbrain | R | 221 | 9.20 | 9 | -30 | -15 |
| Thalamus | R | 75 | 10.42 | 12 | -7 | 3 |
| Thalamus | L | 42 | 9.54 | -5 | -12 | 4 |
| Insula | L | 61 | 9.88 | -41 | 14 | 6 |
| Insula | R | 43 | 9.13 | 46 | 9 | 2 |
| Escapability parametric modulation - : Fast Predator |  |  |  |  |  |  |
| Parahippocampal Gyrus | L | 24 | 4.98 | -21 | -12 | -24 |
| Parahippocampal Gyrus | R | 19 | 4.21 | 39 | -39 | -24 |
| Middle Frontal Gyrus | L | 48 | 7.06 | -9 | 60 | 6 |
| Escapability parametric modulation + : Slow Predator |  |  |  |  |  |  |
| Middle Temporal Gyrus | L | 86 | 5.62 | -57 | -35 | -6 |
| Insula | R | 44 | 6.23 | 39 | -14 | 8 |
| Superior Occipital Gyrus | L | 30 | 4.57 | -14 | -89 | 19 |
| Superior Occipital Gyrus | R | 54 | 4.24 | 17 | -88 | 19 |
| Escapability parametric modulation - : Slow Predator |  |  |  |  |  |  |
| Middle Frontal Gyrus | L | 84 | 7.28 | -30 | 60 | 15 |
| ACC | R | 35 | 8.29 | 18 | 42 | 0 |
| Caudate | L | 14 | 4.34 | -9 | -51 | 27 |
| Caudate | R | 20 | 4.92 | 14 | 15 | -3 |

**Table S5.**
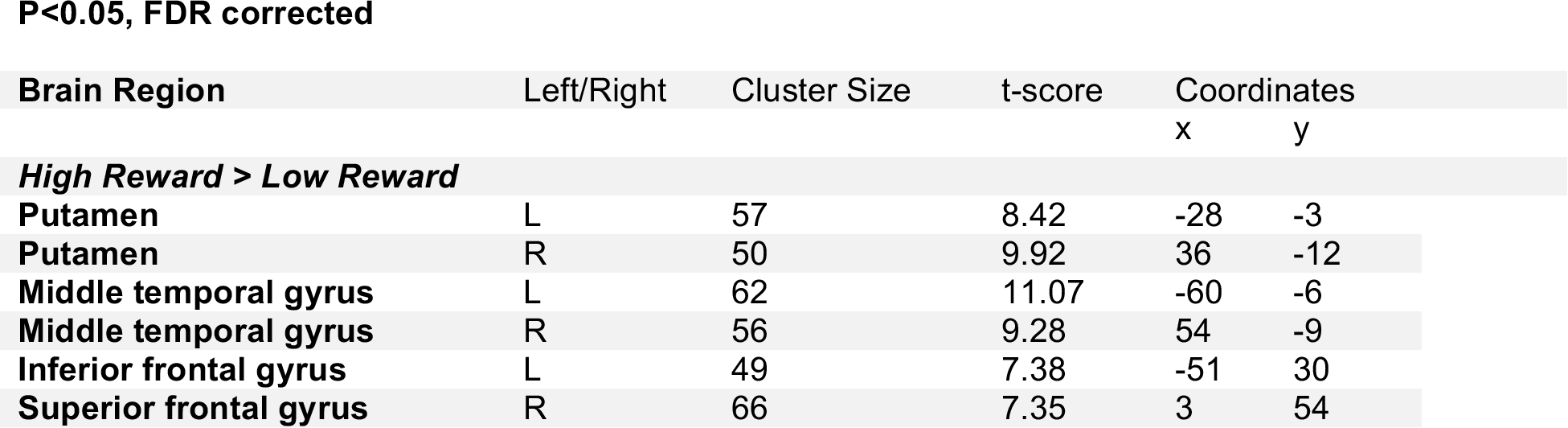
Activation Table for Contrast [High Reward > Low Reward].

**P<0.05, FDR corrected**
| Brain Region | Left/Right | Cluster Size | t-score | Coordinates |  |
| --- | --- | --- | --- | --- | --- |
|  |  |  |  | x | y |
| High Reward > Low Reward |  |  |  |  |  |
| Putamen | L | 57 | 8.42 | -28 | -3 |
| Putamen | R | 50 | 9.92 | 36 | -12 |
| Middle temporal gyrus | L | 62 | 11.07 | -60 | -6 |
| Middle temporal gyrus | R | 56 | 9.28 | 54 | -9 |
| Inferior frontal gyrus | L | 49 | 7.38 | -51 | 30 |
| Superior frontal gyrus | R | 66 | 7.35 | 3 | 54 |

**Table S6.** Activation Table for Contrast [High Shock > Low Shock].

**P<0.05, FDR corrected**
| Brain Region | Left/Right | Cluster Size | t-score | Coordinates |  |  |
| --- | --- | --- | --- | --- | --- | --- |
|  |  |  |  | x | y | z |
| *High Shock > Low Shock* |  |  |  |  |  |  |
| Superior temporal gyrus | L | 147 | 11.16 | -58 | 3 | 6 |
| Insula | L | 54 | 7.36 | -41 | -14 | 3 |
| Insula | R | 35 | 6.04 | 39 | 3 | -12 |
| MCC | R | 32 | 5.96 | 6 | -9 | 39 |
| Parahippocampal gyrus | L | 10 | 9.78 | -24 | -18 | -9 |
| Hippocampus | R | 20 | 7.55 | 36 | -15 | -18 |

**Table S7.** Activation Table for Contrast [High Shock > Low Shock] (Fast Predator)

**P<0.05, FDR corrected**
| Brain Region | Left/Right | Cluster Size | t-score | Coordinates |  |  |
| --- | --- | --- | --- | --- | --- | --- |
|  |  |  |  | x | y | z |
| **High Shock > Low Shock (fast predator)** |  |  |  |  |  |  |
| Midbrain | L | 190 | 4.95 | -1 | -32 | -13 |
| ACC | R | 179 | 4.36 | 3 | 25 | 7 |
| Precuneus | L | 335 | 4.03 | -27 | -54 | 8 |

**Table S8.** Activation Table for Contrast [High Shock > Low Shock] (Slow Predator)

**P<0.05, FDR corrected**
| Brain Region | Left/Right | Cluster Size | t-score | Coordinates |  |  |
| --- | --- | --- | --- | --- | --- | --- |
|  |  |  |  | x | y | z |
| High Shock > Low Shock (slow predator) |  |  |  |  |  |  |
| Medial temporal gyrus | L | 161 | 4.64 | -42 | -33 | 0 |
| Insula | R | 25 | 4.04 | 40 | -16 | 10 |
| Hippocampus | R | 36 | 4.50 | 21 | -30 | -12 |
| Superior temporal gyrus | R | 26 | 4.30 | 45 | -12 | -3 |
| Amygdala | R | 17 | 4.11 | 24 | 4 | -20 |

**Table S9.** Activation Table for Parametric Modulation with Reward Preference.

**P<0.05, FDR corrected**
| Brain Region | Left/Right | Cluster Size | t-score | Coordinates |  |  |
| --- | --- | --- | --- | --- | --- | --- |
|  |  |  |  | x | y | z |
| ***Parametric modulation with reward preference (fast predator)*** |  |  |  |  |  |  |
| Fusiform gyrus | L | 35 | 5.75 | -42 | -6 | -27 |
| Inferior temporal gyrus | R | 23 | 4.93 | 60 | -9 | -18 |
| Insula | L | 80 | 5.04 | -39 | -12 | 6 |
| Putamen | L | 42 | 4.86 | -28 | -18 | 3 |
| Middle temporal gyrus | L | 81 | 4.11 | -60 | -57 | -3 |
| Putamen | R | 78 | 4.88 | 33 | -6 | 6 |
| ***Parametric modulation with reward preference (slow predator)*** |  |  |  |  |  |  |
| Middle occipital gyrus | L | 85 | 5.86 | -36 | -66 | 0 |
| Caudate | R | 50 | 6.70 | 15 | -3 | 15 |
| Calcarine | R | 38 | 4.79 | 18 | -48 | 3 |
| Insula | R | 35 | 4.48 | 33 | 24 | 3 |
| Caudate | L | 46 | 4.88 | -9 | 3 | 9 |

**Table S10.** Activation Table for Parametric Modulation with Shock Avoidance.

**P<0.05, FDR corrected**
| Brain Region | Left/Right | Cluster Size | t-score | Coordinates |  |  |
| --- | --- | --- | --- | --- | --- | --- |
|  |  |  |  | x | y | z |
| Parametric modulation with shock avoidance (fast predator) |  |  |  |  |  |  |
| Inferior temporal gyrus | R | 21 | 4.40 | 51 | -60 | -18 |
| Midbrain | R | 10 | 4.93 | 21 | -18 | -15 |
| Superior temporal gyrus | R | 21 | 4.83 | 60 | -33 | 12 |
| Inferior frontal gyrus | L | 14 | 4.16 | -45 | 42 | 15 |
| Parametric modulation with shock avoidance (slow predator) |  |  |  |  |  |  |
| PCC | L | 23 | 4.92 | -12 | -39 | 6 |
| Thalamus | R | 35 | 6.27 | 0 | -18 | 15 |
| Middle temporal gyrus | L | 45 | 4.69 | -42 | -66 | 15 |
| Superior temporal gyrus | L | 48 | 5.89 | -60 | -45 | 18 |

**Table S11.** Activation Table for Parametric Modulation with Bayesian optimality.

**P<0.05, FDR corrected**
| Brain Region | Left/Right | Cluster Size | t-score | Coordinates |  |  |
| --- | --- | --- | --- | --- | --- | --- |
|  |  |  |  | x | y | z |
| Parametric modulation with Bayesian optimality (fast predator) |  |  |  |  |  |  |
| ACC | R | 24 | 4.31 | 6 | 33 | 21 |
| Superior frontal gyrus | L | 63 | 4.89 | -24 | 9 | 54 |
| Superior motor area | R | 18 | 4.33 | 12 | 9 | 54 |
| Parametric modulation with Bayesian optimality (slow predator) |  |  |  |  |  |  |
| Hippocampus | R | 20 | 4.18 | 33 | -33 | -3 |
| Middle occipital gyrus | R | 12 | 4.40 | 36 | -87 | -3 |
| Precentral gyrus | R | 41 | 4.32 | 63 | -3 | 24 |
| Precentral gyrus | L | 18 | 4.05 | -51 | -3 | 24 |

**Table S12.**
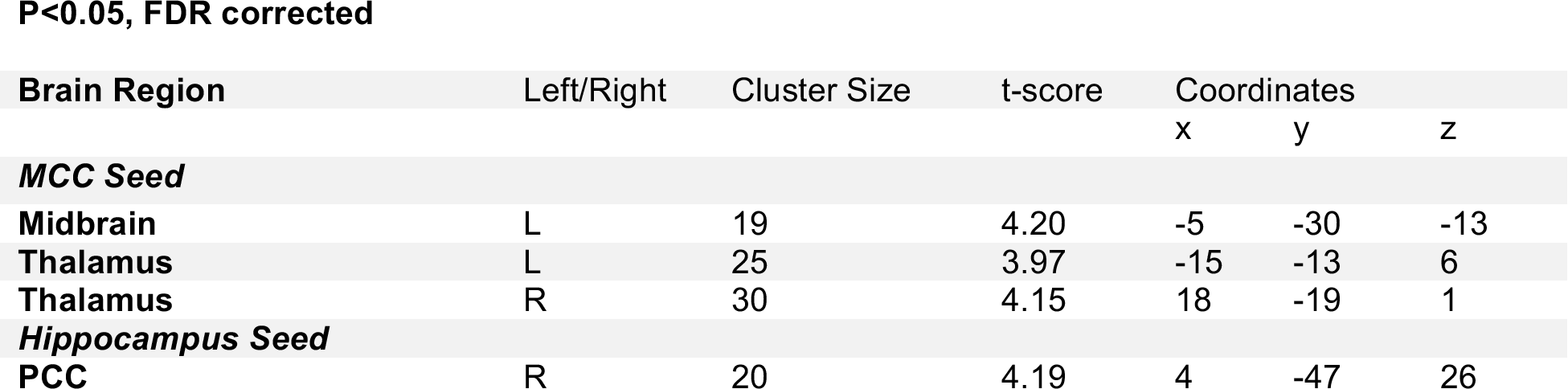
Activation Table for Connectivity Analysis.

